## Supplementary material for "*Drosophila suzukii* avoidance of microbes in oviposition choice": Figure S1: Sato etal-figureS1.pdf

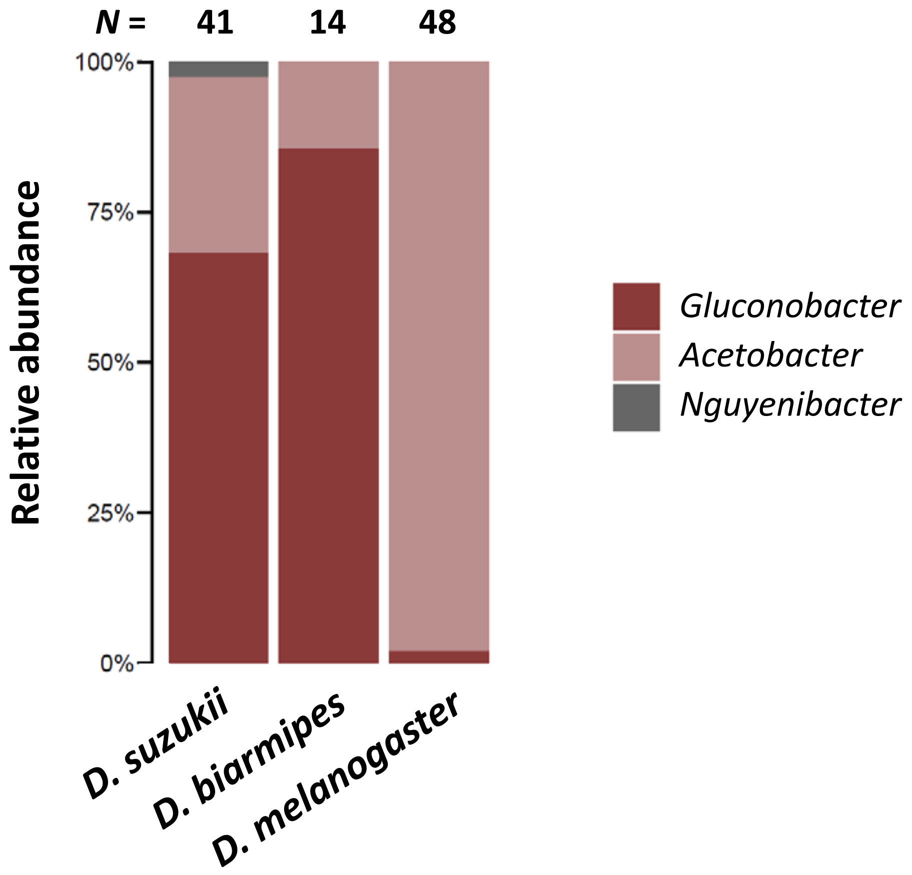


Figure S1. Relative abundance of bacterial taxa grown on 1% agar in 50% apple juice medium after aqueous solutions collected from inoculated media were spread onto new media plates and incubated for 24–40 h. Proportions of bacterial species in each of the three genus identified using 16S rRNA gene sequences from single colonies are indicated in different colors. *Drosophila* species used for inoculation are indicated below the bar graphs. Numbers of sequenced colonies are indicated above the graphs.
